# SiLiCO: A Simulator of Long Read Sequencing in PacBio and Oxford Nanopore

**DOI:** 10.1101/076901

**Authors:** Ethan Alexander García Baker, Sara Goodwin, W. Richard McCombie, Olivia Mendivil Ramos

## Abstract

**Summary:** Long read sequencing platforms, which include the widely used Pacific Biosciences (PacBio) platform and the emerging Oxford Nanopore platform, aim to produce sequence fragments in excess of 15-20 kilobases, and have proved advantageous in the identification of structural variants and easing genome assembly. However, long read sequencing remains relatively expensive and error prone, and failed sequencing runs represent a significant problem for genomics core facilities. To quantitatively assess the underlying mechanics of sequencing failure, it is essential to have highly reproducible and controllable reference data sets to which sequencing results can be compared. Here, we present SiLiCO, the first *in silico* simulation tool to generate standardized sequencing results from both of the leading long read sequencing platforms.

**Availability:** SiLiCO is an open source package written in Python. It is freely available at https://www.github.com/ethanagbaker/SiLiCO under the GNU GPL 3.0 license.

**Contact:** <emails>

**Supplementary information:** Supplementary data are available at *Bioinformatics* online.

## 1 Introduction

Long-read sequencing technology has proved invaluable to overcome the shortcomings of short read sequencing technology. Moreover, the use of long read sequencing has increased exponentially in recent years and continues to do so (Chaisson *et al.* 2015, Goodwin *et al.* 2016). However, quantifying *in silico* the causes of the probabilistically higher rate of sequencing failure in long read sequencing, paired with cost, substantial opportunity remains to improve performance of long read sequencing. It is necessary to have highly reproducible and controllable reference data to which *in vitro* sequencing results can be compared to identify variables implicated in long read sequencing failure.

One observable source of failure in long read sequencing is library performance in the instrument (PacBio and Nanopore) after passing all pertinent molecular quality checks. We hypothesize that NGS libraries that fail sequencing contain an abnormally high degree of nicks, resulting in incomplete sequencing (data not shown). Given that long read sequencing remains relatively expensive in terms of both time and resources, failed sequencing runs represent a significant challenge, particularly for genomics core facilities. Therefore, there is substantial need to understand the underlying mechanics of sequencing failure, including whether some nucleotide combinations are more prone to damage than others. To quantify the patterns of nick sites in sequencing libraries, we developed an *in silico* simulator of both long read sequencing platforms to create an empirical distributions of terminal nucleotides in ideal long read sequencing libraries.

Here, we introduce SiLiCO, the first open source package for *in silico* simulation of long read sequencing results on both major long read sequencing platforms (Sup. Table 1). SiLiCO simulates both PacBio and, for the first time, Oxford Nanopore read sequencing results by stochastically generating genomic coordinates and extracting corresponding nucleotide sequences from a reference assembly. Futhermore, SiLiCO also is easily scaled up to a Monte-Carlo simulation, affording the end user the ability to construct empirical distributions of various genomic features.

## 2 Methods and Implementation

### 2.1 Selection of Read Length Distributions

To approximate the observed distribution of read lengths in both the PacBio and Oxford Nanopore platforms, a model-fitting approach was implemented on exemplary data sets from each platform. For PacBio data, previous literature has determined that PacBio results follow an approximately log-normal distribution with a mean read length of 3kb (Roberts *et al.* 2013). For Oxford Nanopore, published data sets (n=2) from Mikheyev *et al.* 2014 were used and Loman *et al.* 2015. Several candidate distributions (Weibull, gamma, log-normal) were fitted to Oxford Nanopore data sets. While the distribution of Oxford Nanopore read lengths is difficult to model concisely due to a high degree of right-skewedness, it was determined that the gamma distribution best fits Oxford Nanopore data. This determination was made on the basis of a comparison of Akaike information criteria (AIC), a metric that quantifies goodness of fit and information loss of candidate models, and performance of candidate models on Cullen-Frey and probability plots (Figures 1A, 1B, S1, S2, Table S2).

**Figure 1:**
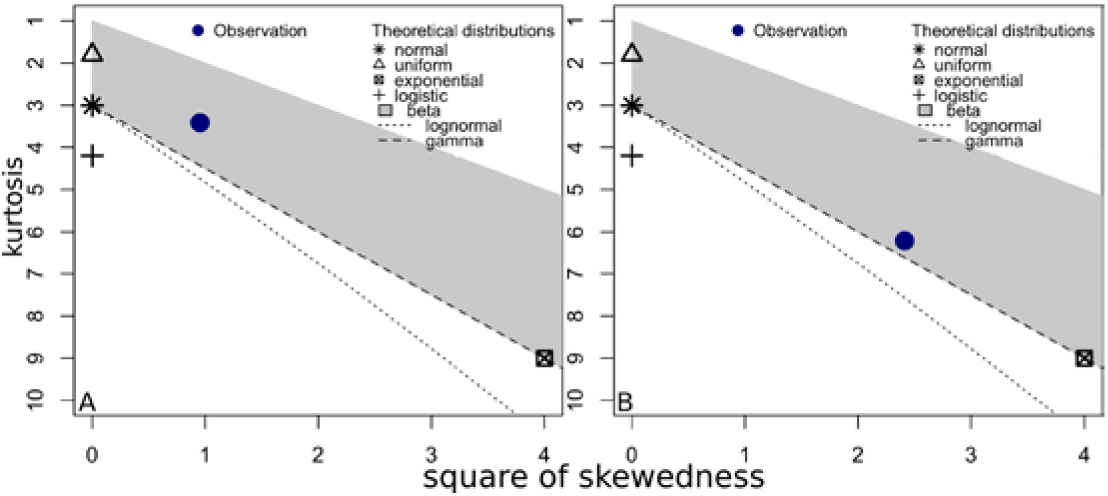
Cullen & Frey plots summarizing the properties of the read-length distribution for both long sequencing platforms PacBio (A) and Oxford Nanopore (B) of the genome sequence of *E. coli.*

### 2.2 *In silico* Read Generation

SiLiCO stochastically generates genomic coordinates based on user-supplied parameters for mean read length, standard deviation of read length, and desired genome coverage. Parameters for mean and standard deviation are mathematically converted to the µ and ∂ parameters of the log-normal distribution and the shape and scale parameters of the gamma distribution when the PacBio and Nanopore modes are invoked, respectively. Read lengths are generated randomly from the user-parametrized distribution, and coordinates are generated. SiLiCO uses the reference genome to ensure that simulated reads lie exclusively on one chromosome. SiLiCO also prevents end-selection bias by selecting start and end coordinates using a buffer, ensuring that all nucleotides have equal likelihood of being selected in a simulated read. SiLiCO executes this algorithm until it achieves the user-specified coverage level, and writes simulated coordinates to a BED file (Figure 2). Optionally, the end user can choose to invoke an implementation of *pybedtools* with the *–-fasta* option to obtain FASTA sequences for simulated reads (Dale *et al.* 2011). Additionally, SiLiCO can be easily scaled to a Monte Carlo simulation using the *--trials* parameter, allowing for simply construction of empirical distributions of genomic features.

**Figure 2:**
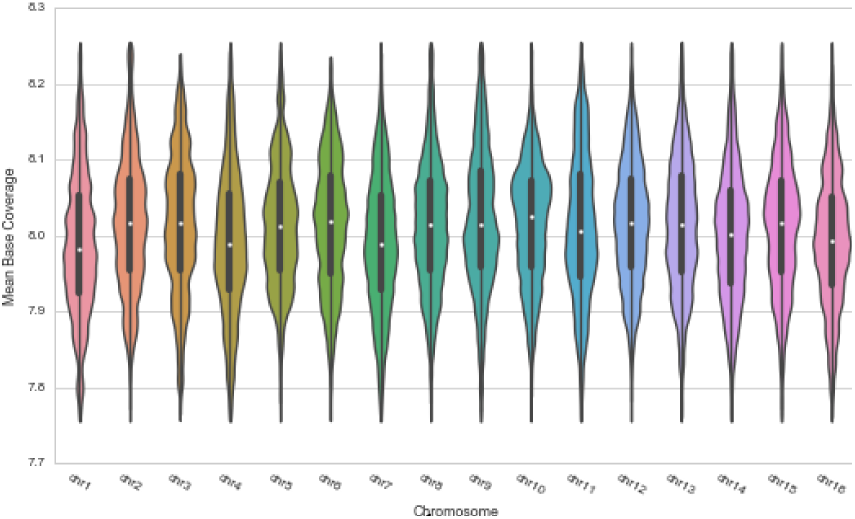
Violin plot of the simulated genome sequence of *Zea mays* produced by SiLiCO. As per user specifications, it produces an even coverage across chromosomes, here shown as the mean base coverage from a Monte-Carlo simulation of PacBio sequencing of *Zea mays* at 8x coverage.

SiLiCO is implemented in Python v. 2.7.11 and relies on numpy for randomization, natsort (https://pypi.python.org/pypi/natsort) for natural sorting, and pybedtools (van der Walt, *et al.* 2011).

## 3 Benchmarking and Validation

A Monte Carlo simulation with 1000 trials of *in silico* sequencing results was generated using SiLiCO. Analysis of simulated results confirmed that SiLiCO produces read lengths consistent with log-normal and gamma distributions and produced even coverage across all chromosomes at the user specified level (Figures S3, S4). Because of SiLiCO’s distribution sampling approach to *in silico* read generation, a reference genome from any species can be used.

SiLiCO was benchmarked on a typical consumer grade computer (Apple MacBook Air with 1.3 GHz Intel Core i5 CPU and 4 GB DDR3 RAM) and can generate a single *in silico* result in ~3.5 seconds and a Monte Carlo simulation of 1000 results in ~8.3 minutes (Sup. Table 3).

## 4 Case Usage

To date and to the authors’ knowledge, no attempts have been made to quantify both the extent of long read sequencing failure and, furthermore, its underlying mechanism. Given our hypothesis that nicking in DNA sequences are implicated in sequencing failure, obtaining data of insufficient quality, we sought to quantify the both the composition and pattern of overrepresentation of nick sites in NGS libraries that fail during sequencing. In order to characterize the differences in nick site composition between healthy and damaged libraries, we compared terminal nucleotide pairs (where sequencing terminated) between libraries that failed long-read sequencing and those that were successfully sequenced with results of adequate quality.

We used SiLiCO to perform a Monte Carlo simulation (n=1000) to determine the empirical distributions of terminal nucleotide pairs in an ideal PacBio sequencing libraries. Terminal nucleotide pairs were selected as a measure of where the modified DNA polymerase employed in PacBio SMRT sequencing stops, possibly at a nick site. *In silico* reads were compared to poorly performing (“failed”) *in vitro* sequencing data to identify and evaluate potential patterns in sites of DNA nicking.

Empirical distributions of terminal nucleotide pairs from *in silico* data were compared with poorly performing *in vitro* PacBio sequencing results to determine patterns of terminal nucleotide pair over/under-representation in long read sequencing failure. Comparisons between *in vitro* and *in silico* libraries prove sufficient to resolve non-stochastic biases in sites of DNA strand breaks. Interestingly, upon replication of this analysis in several species, including *E. coli* K12 and *S. cerevisiae* W303, the observed pattern of over or under representation of nucleotide pairs was largely conserved. We conclude that this conserved pattern suggests preferential occurrence of DNA nicking on certain nucleotide pairs (all individual *p-values* < .01, Z-test for difference of proportions, Figure 3, Table S4).

**Figure 3:**
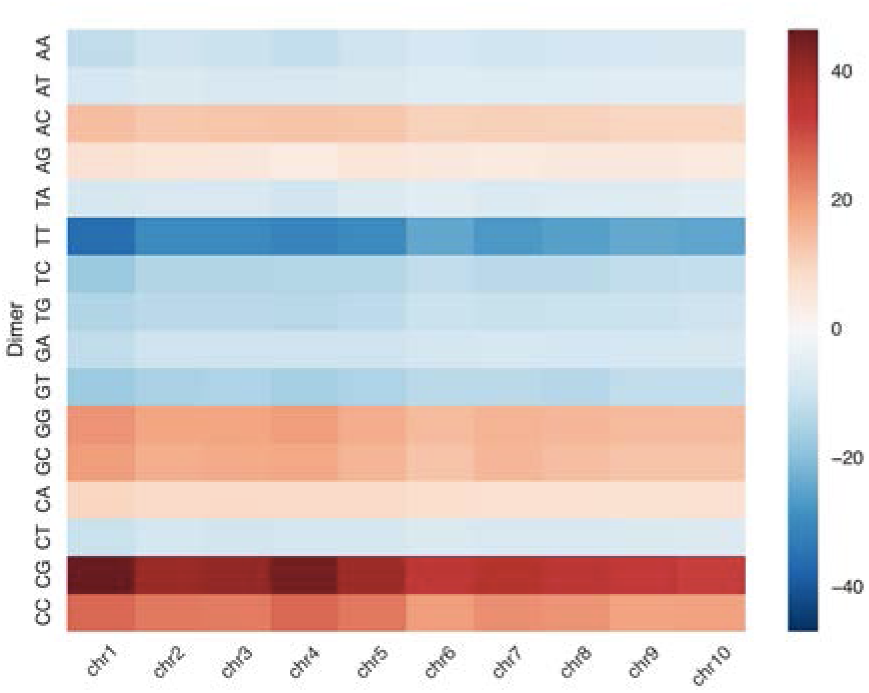
Heatmap of the Z-Score differences in distributions of terminal nucleotide pairs in *Zea mays* between SiLiCO simulated PacBio sequencing and an *in vitro* PacBio library. This library produced abnormally short read lengths, thereby failing to meet normal performance metrics of the PacBio platform. Similar comparisons between *in silico* and *in vitro* sequencing results were used poorly to evaluate patterns in sites of DNA strand breaks.

## Acknowledgements

The authors thank Melissa Kramer, Jayon Lihm, Shane McCarthy, Doreen Ware, the Ware Lab, and the Technology Development Group (McCombie Lab), and Michael Schatz for their comments on this work.

Maize sequencing was performed at the Ware Lab (Yinping Jiao and Bo Wang) at CSHL and supported by NSF Improving Plant Genome Annotation grant 1127112, NSF Cereal Gene Discovery grant 1032105, USDA ARS CRIS 1907-21000-030-00D and NSF Rare Alleles grant 1238014.

This work has been supported by the Undergraduate Research Program at Cold Spring Harbor Laboratory and the National Science Foundation.

